# Larger lesion volume in people with multiple sclerosis is associated with increased transition energies between brain states and decreased entropy of brain activity

**DOI:** 10.1101/2022.02.17.480876

**Authors:** Ceren Tozlu, Sophie Card, Keith Jamison, Susan A. Gauthier, Amy Kuceyeski

**Author notes:** **Corresponding Author:** Amy Kuceyeski, Ph.D., **Mailing address:** 526 Campus Road, Biotechnology building, Room 101D, Ithaca NY 14850, **Email:**.

## Abstract

**Background:** Neuronal loss, demyelination, and inflammation in the brains of people with multiple sclerosis (pwMS) can cause impairment or disability across a range of domains. However, the mapping between severity or location of this brain tissue damage and an individual’s impairment remains challenging. Advanced neuroimaging techniques may enable us to better understand the neuropathological or compensatory mechanisms in MS and quantify the link between the brain’s functional activity and its structural backbone. One recently developed approach uses network control theory to characterize the brain’s energetic landscape using an individual’s structural connectome and brain activity dynamics. While brain activity dynamics have previously been analyzed in MS cohorts, investigating how structural network damage, one of the hallmarks of MS, relates to shifts in activity dynamics has not yet been performed.

**Aims:** We apply brain activity dynamics analysis and network control theory (NCT) to functional time series and structural connectomes in healthy controls (HC) and pwMS with and without disability and compare the three groups’ resulting metrics. We also assessed the entropy of the three groups’ brain activity time series. Our hypotheses were that pwMS with disability would have increased brain state transition energies compared to pwMS without disability and controls, and that greater energies would be related to greater lesion burden and decreased entropy of brain activity.

**Materials and Methods:** Ninety-nine pwMS and 19 HC were included in our study; individuals underwent MRI, including anatomical, diffusion and resting-state functional scans. Expanded Disability Status Scale (EDSS) was used to assess disability; 66 pwMS with EDSS<2 were considered as not having disability. Commonly recurring brain states were identified using k-means clustering of the regional brain activity time series. NCT was used to compute the energy required to transition between each pair of states or to remain in the same state, taking into account an individual’s structural connectome. The entropy of brain activity time series was quantified and correlated with overall transition energy and lesion volume. Finally, logistic regression with ridge regularization was used to classify the groups using transition energies and demographics/clinical information and the model’s feature importance assessed.

**Results:** Six brain states were identified, including pairs of states with high and low visual (VIS+/-), ventral attention (VAN)+/- and somatomotor (SOM+/-) network activity. Global entropy was negatively correlated with lesion volume (r=-0.20, p=0.03) as was global transition energy (r=-0.18, p=0.04) in pwMS. Smaller transition energies were associated with pwMS without disability, while larger transition energies were associated with pwMS with disability.

**Discussion:** We conjecture that functional compensation of brain activity in pwMS without disability results in decreased transition energies between states compared to controls, but, as this mechanism diminishes, transition energies increase and disability occurs. We also conjecture that larger volumes of MS-related lesions result in greater energetic demands to transition between brain states which then leads to lower overall entropy of brain activity.

## Introduction

Multiple Sclerosis (MS) is a chronic disease characterized by neuroinflammation and, eventually, neurodegeneration in the central nervous system (CNS). The size and location of the lesions in the CNS are very heterogeneous^1^, resulting in different patterns of structural damage among people with MS (pwMS). Damage to the brain may result in changes its functional activity patterns^2^, however, how different patterns of structural damage can modify dynamics of functional activity have not been fully characterized in pwMS.

Functional MRI (fMRI) is an important tool with which to measure the brain’s activity patterns over time. Both increased or decreased functional activity in particular brain regions has been observed in pwMS, where increased functional activity/connectivity is often interpreted as a compensatory mechanism limiting clinical disability.^3,4^ FMRI in pwMS has largely been used to investigate the association between the brain’s static and/or dynamic functional connectivity (FC) and motor/cognitive impairment.^5–7^ The majority of FC studies have assumed that the FC is static^3,4^ and have ignored shorter scale changes of brain network activity that have been shown to occur.^8^ The dynamic FC (dFC) approach captures recurrent co-fluctuation states by clustering dynamic FC matrices computed over sliding windows of BOLD time series.^9^ This approach was previously used to differentiate between the pwMS and HC^10,11^, to compare the dynamics between cognitively impaired vs preserved pwMS ^12^, and to investigate the relationship between alterations in the dFC dynamics and cognition in MS.^13–15^ However, calculating dFCs with a sliding window approach is not straightforward since the estimation of the window length and shift size of the sliding windows is required, moreover correlations are still used to describe co-activations of pairs of regions as in static FC. As an alternative approach, clustering can be directly applied to the time series of regional activity to define distinct brain states. This approach was successfully used in healthy controls (HC) to identify dynamic brain states that occur while performing a working memory task as well as quantify the effects of psychedelics.^16,17^ In addition to the identification of the brain dynamic states, the same studies used the network control theory (NCT) approach^18–20^ to identify the minimum energy required to transition between these dynamic brain states. However, no study to date has applied brain activity clustering and NCT in a population of pwMS, let alone investigate differences in brain dynamics and energetics across disability subgroups in MS.

Entropy of brain activity over time is a fundamental metric to quantify the amount of regularity/unpredictability in BOLD time series.^21^ Higher entropy of brain activity has been associated with a larger repertoire of available brain states^22^ and the amount of decrease in transition energy under psychedelics (compared to placebo) was found to be associated with greater increases in entropy of brain states.^17^ Other studies have shown that entropy is associated with lesion size and disability level in stroke and attention deficit hyperactivity disorder (ADHD) or MS.^23–25^ However, no study to date has investigated the link between the entropy of brain activity and 1) lesion volume or 2) transition energy between brain states in MS.

In this study, we first aim to identify recurrent dynamic brain states in both HC and pwMS by unsupervised clustering of regional resting-state fMRI activity. We then calculate metrics quantifying the dynamics of these states, including transition probability and fractional occupancy, and use NCT and individuals’ structural connectomes (SC) to compute the energy required to transition between pairs of brain states. We hypothesize that greater energy is required for brain state transitions in pwMS with disability compared to both controls and pwMS with no evidence of disability. Finally, we calculated the entropy of each region’s activity over the entire fMRI scan and summarize the global entropy by taking the average of the regional measures. Our hypothesis is that decreased global entropy is associated with both increased lesion load and transition energy. This paper is the first to quantify MS-related shifts in the brain’s energetic landscape and to link entropy/energetic demand of brain activity to lesion burden in pwMS.

## Material and Methods

### Subjects

One hundred pwMS (age: 45.5 [36.7, 56.0], 66% females) with a diagnosis of CIS/MS (CIS=7, RRMS=88, primary/secondary progressive PPMS and SPMS = 5) and 19 HC (age: 45 [35, 49], 55% females) were included in our study. MRIs and demographic data were collected (age, sex, and race for both HC and pwMS, clinical phenotype and disease duration for pwMS). EDSS was used to quantify disability in pwMS, where an EDSS of 2 was used as a threshold to categorize the pwMS into no disability (EDSS<2) or evidence of disability (EDSS≥2) groups. This classification was based on EDSS values of 0-1.5 representing some abnormal signs but no functional disability appreciated. All studies were approved by an ethical standards committee on human experimentation and written informed consent was obtained from all patients. Participants were excluded if they had contraindications to MRI and controls were further excluded if they had ever been diagnosed with or were currently on medication for a neurological or psychological disorder.

### Image acquisition, processing, and structural connectome extraction

MRI data were acquired on a 3T Siemens Skyra scanner (Siemens, Erlangen, Germany) with a 20-channel head-neck coil and a 32-channel spine-array coil. Anatomical MRI (T1/T2/T2-FLAIR, 1 mm3 iso-voxel), resting-state fMRI (6 min, TR = 2.3 s, 3.75 x 3.75 x 4 mm voxels) and diffusion MRI (55 directions HARDI, b=800, 1.8 x 1.8 x 2.5 mm voxels) acquisitions were performed. Multi-echo 2D GRE fieldmaps were collected for use with both fMRI and diffusion MRI (0.75 x 0.75 x 2 mm voxels, TE1=6.69 ms, ΔTE=4.06 ms, number of TEs = 6).

White matter (WM) and gray matter (GM) were segmented and GM further parcellated into 86 FreeSurfer-based regions (68 cortical and 18 subcortex/cerebellum)^26^ and a fine-grained atlas cc400 (321 cortical and 71 subcortex/cerebellum) using FreeSurfer ^27^. As described elsewhere^28^, fMRI preprocessing included simultaneous nuisance regression and removal of WM and cerebrospinal fluid (CSF) effects^29^, followed by high-pass filtering (≥0.008) using the CONN v18b toolbox^30^ and SPM12 in Matlab. Nuisance regressors included 24 motion parameters (6 rotation and translation, temporal derivatives, and squared version of each) and the top 5 eigenvectors from eroded masks of both WM and cerebro-spinal fluid. The mean fMRI signal over all voxels in each of the 86 regions for each TR was calculated to obtain a regional time series matrix of BOLD activity.

Diffusion MRI was interpolated to isotropic 1.8mm voxels, and then corrected for eddy current, motion, and EPI-distortion with the eddy command from FSL 5.0.11^31^ using the outlier detection and replacement option.^32^ MRtrix3Tissue (https://3Tissue.github.io), a fork of MRtrix3^33^ was used to estimate a voxel-wise single-shell, 3-tissue constrained spherical deconvolution model (SS3T-CSD) and then compute whole-brain tractography for each subject. We performed deterministic (sd-stream) tractography with MRtrix3^34,35^ with uniform seeding at each white-matter voxel, after which the SIFT2 global filtering algorithm ^36^ was applied to account for bias that exists in greedy, locally-optimal streamline reconstruction algorithms. The SC matrix was constructed by taking the sum of the SIFT2 weights of streamlines connecting pairs of regions and then dividing by the sum of the two regions’ volumes.

Lesion masks were produced for each pwMS in a semi-automated process. The WM hyperintensity lesion masks were created by running the T2 FLAIR images through the Lesion Segmentation Tool (LST) and were further hand-edited as needed. T2 FLAIR-based lesion masks were transformed to the individual’s T1 native space using the inverse of the T1 to GRE transform and nearest-neighbor interpolation. Individual T1 images were then normalized to MNI space using FSL’s linear (FLIRT) and non-linear (FNIRT) transformation tools (http://www.fmrib.ox.ac.uk/fsl/index.html); transformations with nearest-neighbor interpolation were then applied to transform the native anatomical space T2FLAIR lesion masks to MNI space. The transformations were concatenated (T2FLAIR to T1 to MNI) to minimize interpolation effects. Lesions were manually inspected after the transformation to MNI space to verify the accuracy of coregistration and lesion volume (in mm^3^) calculated.

### Brain state identification

K-means clustering was used to identify commonly recurring brain states (see Figure 1 for the workflow of the study) over all subjects’ regional brain activity time series. We used both the elbow criterion and the gain in explained variance to identify the optimal number of clusters, which we identified to be 6 (see Supplementary Figure 2). Once the optimal number of clusters was identified, clustering was run 10 times. Within each of the 10 runs, clustering was restarted 50 times with different random initializations to identify the final clusters having the best separation, i.e. those that minimized the Pearson correlation between pairs of cluster centroids. Adjusted mutual information (AMI) ^37^ was computed to assess the stability of the clustering results over the 10 repetitions; AMI was greater than 0.89 for all pairs of repetitions indicating robust cluster assignment (see Supplementary Figure 1).

**Figure 1:**
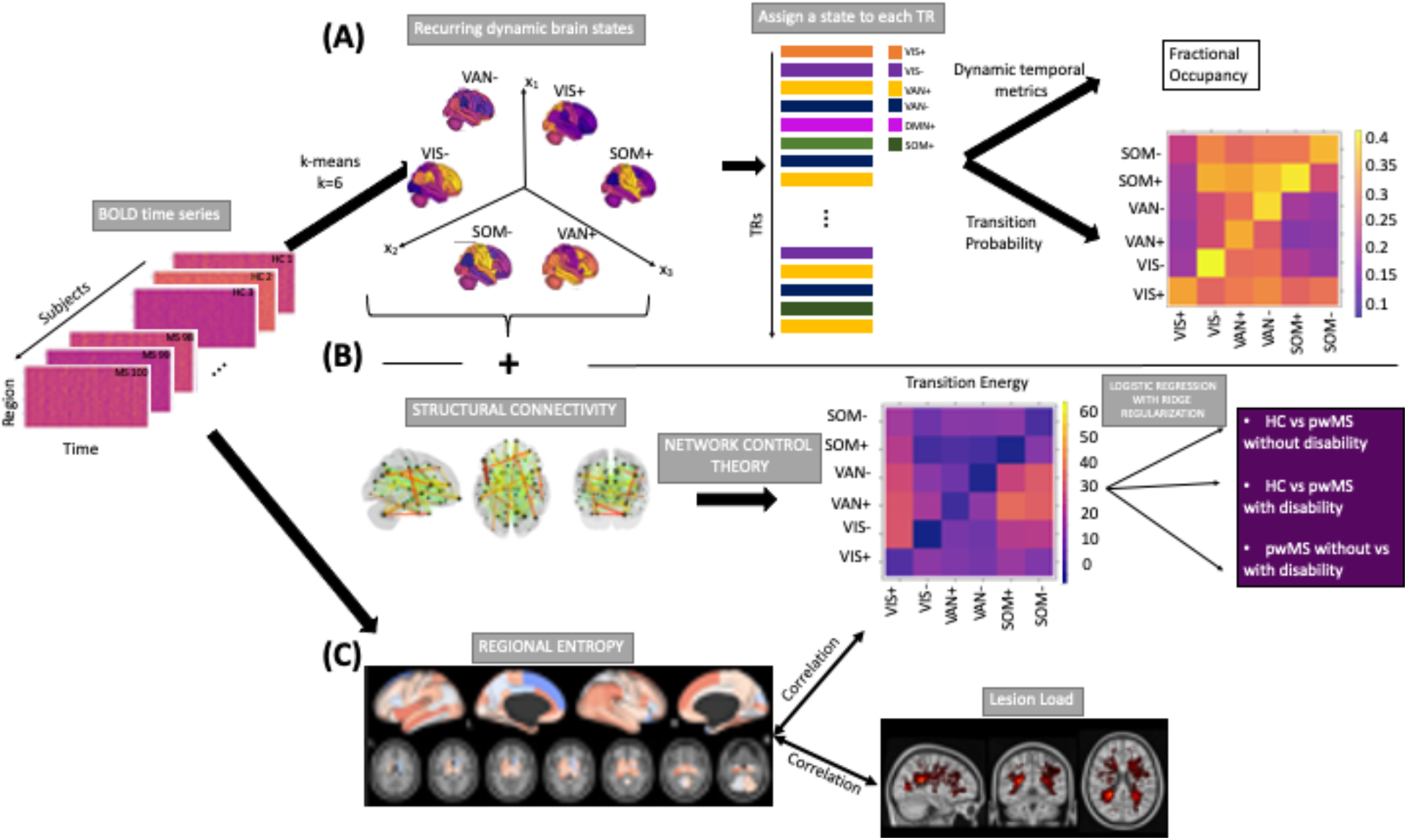
The workflow of the study. (A) Regional BOLD time series for all subjects (19 HC and 99 pwMS) were collected and k-means was applied to identify commonly occurring brain states. This approach allows us to assign a state to each TR in each individual’s scan. Metrics such as fractional occupancy and transition probability for each state were then calculated. (B) Network control theory was applied to compute the minimum energy required to transition between pairs of brain states or to remain in the same state. Logistic regression with ridge regularization was used to predict class (HC, pwMS with no disability, pwMS with disability) using state transition/persistence energy (C) Entropy of regional BOLD time series were also computed, compared across the three groups and global entropy was associated with global transition energy and global disease burden (log lesion volume).

Finally, we replicated the cluster analysis using a higher dimensional atlas (cc400), where optimal k was identified as 5 (See Supplementary Figure 6). The results obtained with the cc400 atlas were interpreted in the main text and the figures associated with those results were given in the supplementary materials.

Each of the 86 regions was assigned to one of 9 networks, including the 7 Yeo functional networks, a subcortical and a cerebellar network.^38^ The network-level contributions of the six centroids were characterized by calculating the cosine similarity of each state’s centroid (positive and negative parts were analyzed separately) and 9 binarized vectors representing each of the networks. Because the mean signal from each scan’s regional time series was removed during high-pass filtering, positive values reflect signal intensity above the mean (high-amplitude) and negative values reflect signal intensity below the mean (low-amplitude). The brain states were named by identifying the network with the largest magnitude activation (either high+ or low- amplitude). The following dynamic brain state metrics were calculated: 1) transition probability between pairs of states and the persistence probability of remaining in the same state and 2) fractional occupancy, defined as the number of TRs assigned to each cluster out of the total number of TRs.

### Transition energy calculations

As previously described^16,17^, NCT can be used to understand how the structural (white matter) connectome constrains dynamic brain state changes. Specifically, it can be used to compute the minimum energy (*E_m_*) required to transition between pairs of brain states or to remain in the same state. To begin, we employed a time-invariant model:

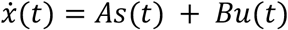

where *A* is the structural connectivity matrix with a dimension of 86×86, *s(t)* is a vector of length 86 containing the regional activity at time *t*, and *u(t)* is the external input to the system. In our case, *B* is an identity matrix with a dimension of *86×86* that indicates each region has uniform control over the system.

When computing the transition energy (*E_m_*) from the initial activity *s*_0_ to final activity *s_f_* over some time *T*, we use an invertible controllability Gramian *W* to control the structural connectivity network *A* from the set of network nodes *B*. The invertible controllability Gramian *W* is defined as:

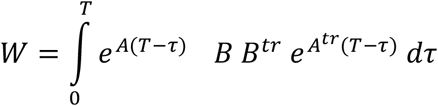

Optimal *T* was identified by grid searching between 0.001 and 10 and identifying the value that maximized the magnitude of the Spearman correlation between the entries in the transition probability matrix and the entries in the transition energy matrix. The correlation between transition probability and energy is expected to be negative since the brain prefers trajectories through state space requiring minimal input energy given structural constraints, therefore the brain tends to change less between states if the minimum required energy is greater. Here, we identified an optimal *T* of 1, which yielded a correlation of −0.60 (p-value<0.001) (see Supplementary Figure 5), while the optimal *T* was 0.5 for the cc400 atlas with a correlation coefficient of −0.90 (see Supplementary Figure 11).

After computing the invertible controllability Gramian *W* and optimizing *T*, the transition energy (*E_m_*) is then calculated as the quadratic product between the inverted controllability Gramian *W* and the difference between *e^AT^s*_0_, which represents the activation at time *T* assuming brain activity spreads in a network diffusion manner over the SC from the initial state *s*_0_, and the final state *s_T_*:

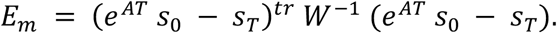

Since there were 6 brain states, the transition energy matrix is of size *6×6* where off-diagonal entries contain the energy required to transition between each pair of unique states *s*_0_ and *s_T_*, where *s*_0_ ≠ *s_T_*, and diagonal entries contain persistence energy required to stay in the same state (*s*_0_=*s_T_*).

### Entropy of brain activity

To investigate how disease burden (lesion load) and disability level in pwMS were related to the amount of information contained in the brain activity signal, regional entropy was calculated directly on the BOLD time series. Denoting the regional BOLD time series in a region by *s*(*t*) = (*s*(1),…,*s*(*n*)) where *n* is the number of the BOLD time series acquisition. We define two template vectors of length m such as: *s_m,i_* = (*s*(*i*), *s*(*i* + 1), …,*s*(*i* + *m* – 1)) and *s_m,j_* = (*s*(*j*),*s*(*j* + 1),…,*s*(*j* + *m* – 1)).Entropy was calculated as the negative logarithm of the probability that if two template vectors of length *m* have distance less than *r* (i.e. *d*[*s_m,i_, s_m,j_*] < *r*) then two template vectors of length *m*+1 also have distance less than *r* (i.e. *d*[*s_m+1,i_*, *s*_*m*+1,*j*_] < *r*) such as

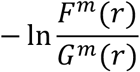

where *F^m^*(*r*) is the number of template vector pairs having *d*[*s*_*m*+l,*i*_, *s*_*m*+1,*j*_] < *r, G^m^*(*r*) is the number of template vector pairs having *d*[*s_m,i_, s_m,j_*] < *r* and where d was the Euclidean distance ^39^. In our study, m = 3, as suggested in two previous studies that computed entropy from restingstate fMRI data in controls^40^ and in pwMS^25^. The parameter r was defined as 0.2 *sd*(*s*) where *sd*(*s*) is the standard deviation of the time series (*s*), as recommended by previous work.^41^ An individual’s global entropy was calculated as the average of their regional entropies.

### Mass univariate analysis

First, demographics and clinical variables were tested for differences between the three groups (HC, pwMS who had no disability, and pwMS who had disability) using Chi-squared test for qualitative variables and Student’s t-test test for quantitative variables. Second, fractional occupancy of and transition probabilities between brain states and regional entropy were compared between groups via Student’s t-tests. Group differences were considered significant when p<0.05 after Benjamini-Hochberg correction for multiple comparisons. Finally, the association between average regional (global) entropy and average (global) transition energy was computed using Spearman’s rank correlation, while the association between global entropy and log of lesion volume (transformed to ensure normality) was computed using Pearson’s correlation. We hypothesized that decreased global entropy would be related to increased disease burden (larger lesion volumes) and increased global transition energies. All statistical analyses and graphs were performed using R (https://www.r-project.org), version 3.4.4 and Matlab (https://www.mathworks.com/) version R2020a.

### Classification analysis

Logistic regression with ridge regularization was used to classify 1) HC and pwMS without disability, 2) HC vs pwMS with disability, and 3) pwMS without vs with disability, using demographics/clinical information (age for HC vs pwMS classification and age and disease duration only for the classification of pwMS subgroups) and vectorized transition energy matrices.

The models were trained with outer and inner loops of k-fold cross-validation (k = 5) to optimize the hyperparameter (λ) and test model performance. The folds for both inner and outer loops were stratified to ensure that each fold contained the same proportion of subjects in the two classes as the original dataset. The inner loop (repeated over 5 different partitions of the training dataset only) optimized the set of hyperparameters that maximized the validation set AUC. A model was then fitted using the entire training dataset and the optimal hyperparameters, which was then assessed on the hold-out test set from the outer loop. The outer loop was repeated for 100 different random partitions of the data. The median AUC (over all 5 folds x 100 iterations = 500 test sets) was calculated to assess the performance of the models.

When the data contains class imbalance, models tend to favor the majority class. Due to the class imbalance in our data (66 vs 33 pwMS with no disability vs evidence of disability), the over-sampling approach Synthetic Majority Over-sampling Technique (SMOTE) ^42^ was used to obtain a balanced training dataset during cross-validation. SMOTE compensates for imbalanced classes by creating synthetic examples using nearest neighbor information and has been shown to be among the most robust and accurate methods with which to control for imbalanced data.^43^

The interpretation of the parameter coefficients from linear models is difficult when there are co-linearities in the data.^44^ To mitigate this problem, we applied the Haufe transform to the model coefficients estimated in each outer loop. Specifically, solving logistic regression with ridge regularization yields model coefficient estimates

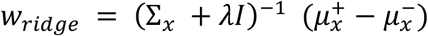

where Σ_*x*_ is the covariance matrix of the input variables *x* in the training dataset, *λ* is the hyperparameter of the ridge classifier optimized in the inner loop for the training dataset, *I* is the identity matrix with the same dimension as 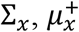 and, 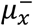 are the average of the input variables for the positive and negative classes in the training dataset. The Haufe transform for these model coefficients is thus

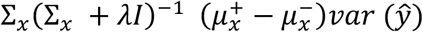

where *ŷ* is the output of the model for the training dataset. The Haufe transformed model coefficients were averaged over all 5 folds x 100 iterations = 500 training sets and the result was used to interpret the relationship between TE and the three groups of individuals.

### Data/code availability statement

The de-identified data that support the findings of this study are available upon reasonable request from the corresponding author. The codes that were used to generate the results and figures are publicly available. Please see 1) https://github.com/cerent/MS-NCT/ for the codes that were used for the classification analysis and figures and 2) the paper of Cornblath et al.^16^ for the codes that were used for the clustering and network control theory approach.

## Results

### Patient Characteristics

Table 1 shows the subjects’ demographic and clinical information including sex, age, disease duration, EDSS, MS phenotype, and lesion volume. Unsurprisingly, pwMS with disability were older than both HCs and pwMS without disability (corrected p < 0.05) and had a trend toward shorter disease duration compared to pwMS without disability (corrected p = 0.15). Sex was not different across any of the groups. Phenotype and disability groups were not independent (corrected p < 0.05), where CIS individuals were included in the without disability group and PPMS individuals were included in the with disability group. The MS subgroups did not have a significant difference in lesion volume (corrected p-value > 0.05).

**Table 1:** Subject demographics and clinical information. Values are presented as median [1st, 3rd quantile] for the continuous variables and as number (percent) for sex. The HC vs pwMS without and with disability as well as the two groups of pwMS were tested for differences; p-values shown are corrected for multiple comparisons, * indicates significance.

| **Variable** | **HC (n=19)** | **pwMS without disability (n=66)** | **pwMS with disability (n=33)** | **p-value (pwMS without disability vs pwMS with disability)** |
| --- | --- | --- | --- | --- |
| **Age** | 45 [35.55, 49.50] | 40.50 [35, 50]<br>p-value vs HC = 0.87 | 56 [46, 58]<br>p-value vs HC = 0.001* | 0.0001* |
| **Female (%)** | 11 (55%) | 46 (70%)<br>p-value vs HC = 0.41 | 20 (61%)<br>p-value vs HC = 0.96 | 0.49 |
| **Disease Duration** |  | 10 [7,14.75] | 13 [9,17] | 0.15 |
| **EDSS** |  | 0 [0, 1] | 2 [2, 3] | <2.2e-16* |
| **Phenotype** |  | 7 CIS, 59 RRMS | 28 RRMS, 5 PPMS | < 0.001* |
| **Lesion Volume (mm<sup>3</sup>)** |  | 1904 [726,4111] | 2482 [453, 7788] | 0.14 |

### Brain states and comparison of dynamics

Figure 2 shows the six brain states, consisting of three pairs of anticorrelated states with high/low amplitude activity in the visual network (VIS+/-), ventral attention network (VAN+/-), and somatomotor network (SOM+/-). The transition probability between dynamic states was not significantly different between HC and any MS subgroup or between MS subgroups (corrected p-value>0.05) (see Supplementary Figure 3). The pwMS subgroups had a trend toward shorter fractional occupancy in the SOM+ and longer fractional occupancy in the VAN+/- and SOM- compared to HC. The pwMS with evidence of disability had a trend toward greater fractional occupancy in the VIS+/-, VAN+, and SOM+ compared to those without disability. However, the differences were not significant for all comparisons (corrected p-value> 0.05) (See Supplementary Figure 4).

**Figure 2:**
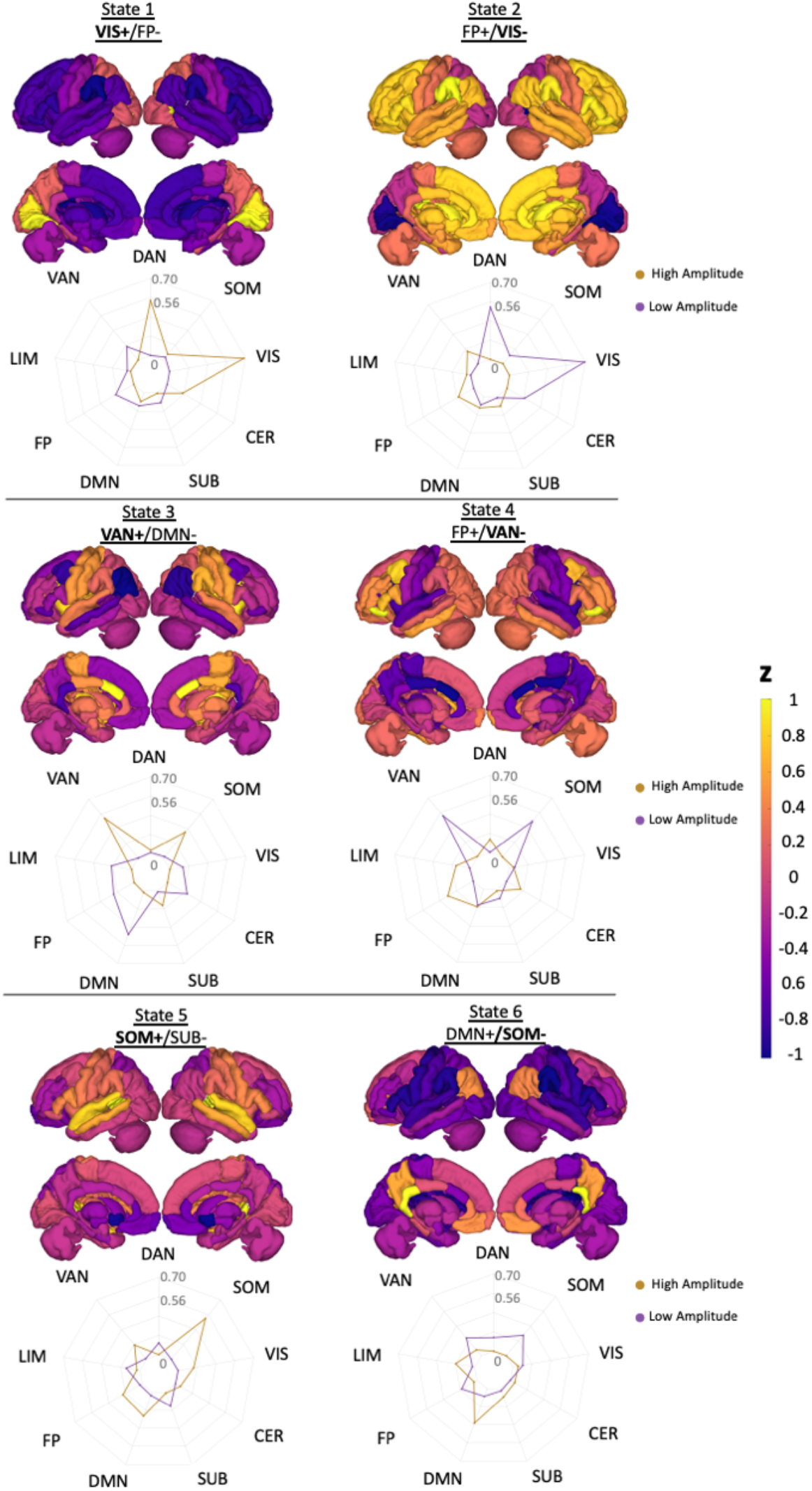
Brain states are visualized using glass brain and radial plots. The radial plots show the mean of the positive and negative brain state values over the Yeo functional networks (DAN= Dorsal Attention, VAN=Ventral Attention, LIM=Limbic, FP=Fronto-Parietal, DMN= Default-Mode Network, SUB=Subcortex, CER=Cerebellum, VIS=Visual, and SOM=Somatomotor). States were named based on the network having the maximum magnitude in the radial plot.

### Entropy of brain activity

Figure 3 shows the t-statistics comparing regional entropy between HC and both MS subgroups and between MS subgroups. There was no significant difference in regional entropy between HC and either MS subgroup (corrected p-value>0.05). However, entropy in the right insula, which is in the VAN network, was significantly decreased in the pwMS with disability compared to those without disability (corrected p-value=0.003). Figure 3 also shows the scatter plot of global entropy and i) global transition energy (average across all entries in the transition energy matrix) and ii) log of lesion volume. Lower entropy was significantly correlated with greater transition energy (r=-0.18, p-value=0.04) and greater lesion volume (r=-0.20, p-value=0.03).

**Figure 3:**
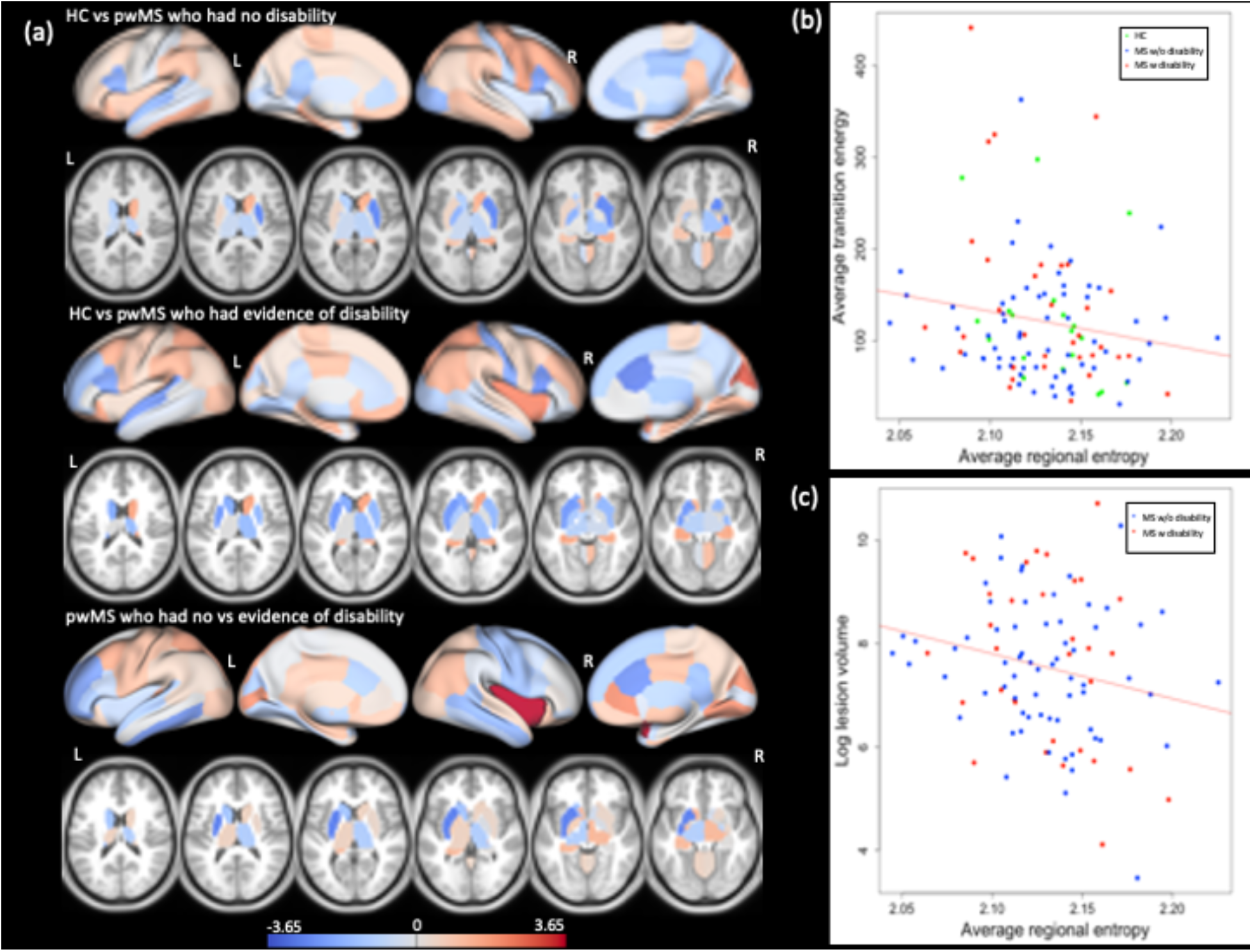
Group difference in entropy and correlation between entropy vs transition energy and lesion load. (a) Group differences in regional entropy between HC vs pwMS without disability (top panel), HC vs pwMS with evidence of disability (middle panel), and between two MS subgroups (bottom panel). The t-statistics presented on the figure indicate HC - MS subgroups and pwMS with no disability - evidence of disability. The scatterplot of global entropy vs (a) global transition energy and (b) log of lesion volume (in mm^3^). The entropy and energy were negatively correlated with average transition energy (r=-0.18, p-value=0.04) and lesion volume (r=-0.20, p-value=0.03). Green points represent HC (only present in the global energy plot), blue dots represent pwMS without disability and red dots represent the pwMs with evidence of disability.

### Group classification using state transition energies

Figure 4 shows the Haufe transformed logistic regression model coefficients for classifying HC vs pwMS without disability, HC vs pwMS with disability, and pwMS without vs pwMS with disability; the models had an average AUC of 0.63, 0.62, and 0.62, respectively. Overall, greater transition energy was associated with pwMS with disability compared to both HC and pwMS without disability, while lower transition energy was associated with pwMS without disability compared to HC. Greater transition energies to and from SOM+ and VIS- and SOM- and VAN+ were associated with pwMS with disability compared to HC, while larger decreases in energy to transition out of VAN+/- was associated with pwMS without disability compared to HC. Greater transition energy out of VAN+ and persistence energy in VAN+ was associated with disability in pwMS compared to those without disability.

**Figure 4:**
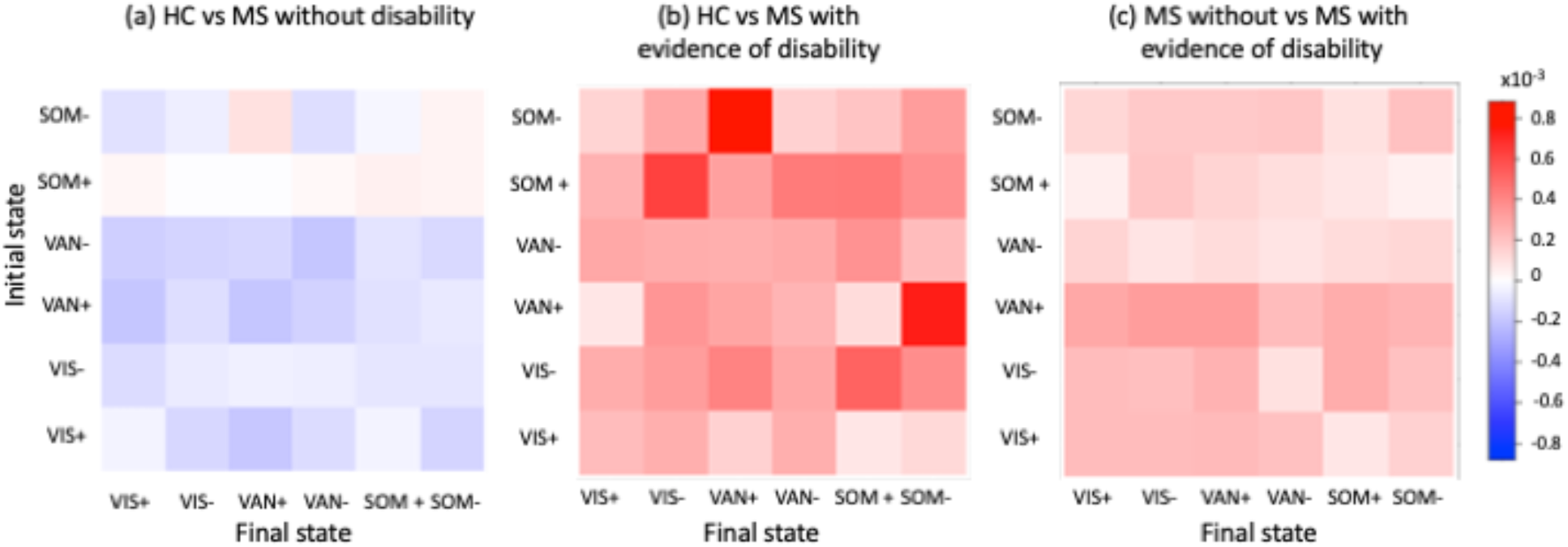
Assessing the relationship between transition energy and disability. Haufe-transformed coefficients for the logistic regression with ridge regularization models classifying HC vs pwMS without disability, HC vs pwMS with disability, and pwMS without disability vs pwMS with disability. (VIS= Visual, VAN= Ventral attention network, and SOM= Somatomotor).

### Robustness of results

The analysis was replicated using the cc400 atlas; five optimal states were found: VIS+/−, SOM+/−, and VAN- (see Supplementary Figure 7), which were overlapping with 5/6 states found with the 86 region atlas. Similar to the original results, there was no significant difference in fractional occupancy between groups (corrected p-value>0.05) (see Supplementary Figure 8). There was a significant negative correlation between global transition energy and global entropy (r=-0.23, p-value=0.01), while the correlation between global entropy and log lesion load was not significant. There were similar results obtained with the 86-region atlas: 1) greater overall transition energy was associated with pwMS who had no disability compared to HC and evidence of disability in MS, 2) greater transition energies between SOM- and VAN- were associated with pwMS who had disability compared to HC.

## Discussion

In this study, we identified commonly occurring brain states in a group of HC and pwMS individuals, investigated the energy required to transition between them using the NCT approach, and, furthermore, associated these energetic measurements with lesion load and brain activity entropy. Our main findings are (1) there were no group differences (HC vs pwMS) in the dynamics of the six recurring brain states, made up of high and low amplitude activity in visual, ventral attention, and somatomotor networks, (2) lower entropy of global brain activity was associated with greater lesion load and global transition energy and (3) pwMS without disability had decreased transition energies compared to controls while pwMS with disability had increased transition energy compared to both controls and pwMS without disability. We hypothesize that this latter finding may indicate a functional compensation mechanism in the pwMS without disability that decreases transition energies, but is eventually exhausted and transition energies increase and disability occurs.

A recent study showed HCs had greater brain activity entropy than people with ADHD, and, further, that lower entropy in people with ADHD was related to worse symptom scores.^23^ Decreased entropy was found in the activity of regions containing a stroke lesion and their contralesional hemisphere homologues.^24^ Our findings are consistent with these previous findings in that increased lesion volume, and likely a weaker structural backbone that may contribute to increased state transition energy, relates to a decrease in the entropy of brain activity. One study found decreased entropy in the right precentral and left parahippocampus and increased entropy in the left superior temporal, right postcentral, and right transverse temporal in RRMS compared to HC compared to controls, which is largely consistent with our regional results ^25^. However, the same study showed no significant correlation between overall lesion volume and the averaged entropy computed for each voxel. The difference in our global findings could be due to a number of factors, including differences in the pwMS cohort characteristics, MRI acquisition and processing differences and differences in the calculation of entropy (voxel-wise as opposed to region-wise entropy in our study). Finally, greater reductions in global transition energy from the placebo to psychedelic state (in controls) were shown to be associated with larger increases in brain state entropy^17^, which is consistent with our findings of negative correlation between global entropy and transition energy across HC and pwMS.

The brain dynamics have previously been studied by clustering regional fMRI time series directly or sliding-window dynamic functional connectivity in HC^16,45,46^ and in patient populations such as stroke^47,48^ schizophrenia^49–51^, and MS^6,11,12^. While many studies investigated the dynamics of sliding-window functional connectivity networks in MS^6,11,12^, this approach has some limitations as the window length and shift size need to be selected in a somewhat ad hoc manner. In this study, we use an alternative approach to identify the dynamic brain states in pwMS, for the first time, by clustering the regional time series activity directly. This approach has an advantage compared to other techniques as it is easy to implement, no ad-hoc parameter estimation is required, and it uses each of the TRs as an individual data point which is especially helpful with short scans where dFC/FC estimates may be unreliable. Using this NCT approach also allow us to investigate the minimum control energy required to transition between dynamic states while incorporating SC, a measure that can capture the topological effects of MS pathology directly.

It has been shown in early MS there may be an upregulation of functional activity that initially acts to limit disability, followed by exhaustion of this functional compensation mechanism as disability increases ^2^. Previous studies showed these adaptive changes might be most prominent in motor and cognition-related networks, thus limiting clinical measures of disability ^52^ as well as cognitive impairment ^53^ in MS. Moreover, the extent of the compensatory mechanism was found to be negatively correlated with the degree of tissue damage^53,54^, such that more damage was related to decreased compensation. Mirroring this finding, our study showed overall lower energetic demand in pwMS without disability compared to controls, which could be reflective of functional upregulation, and a higher energetic demand in pwMS with disability compared to controls, which could be reflective of an exhaustion of functional upregulation and/or increased damage to SCs.

The SOM and VAN states appeared to be the most important in the three classification models, with largest decreases in transition energy out of VAN+/− for pwMS without disability compared to controls and largest decreases in transition energies out of VAN+ for pwMS without disability versus those with disability. The biggest contributions to the HC vs pwMS with disability classification task were transitions to and from SOM- and VAN+. Our recent study showed that greater functional connectivity in the SOM and VAN states was associated with pwMS who had no disability compared to those with disability.^5^ In our current study, the SOM-state also had the highest amplitude DMN activity. The VAN and DMN are known to be highly anticorrelated since the DMN is task-negative and the VAN is a task-positive network. Decreased anticorrelation between DMN and attention networks was related to worse behavioral performance in HC^55^, moreover increased functional connectivity between VAN and DMN was observed in pwMS who had cognitive impairment compared to those without cognitive impairment and HC.^56^ Even though we did not have cognitive scores for our cohort, previous studies have identified a strong association between disability and cognitive impairment in MS.^57^ Our results suggest that dysfunction of the interaction between the VAN and DMN states may result in greater energetic demand between these states in pwMS with evidence of disability compared to the other groups. Finally, our results also showed greater transition energy between SOM+ and VIS-was associated with pwMS who had disability. The most frequently observed disturbances are the motor and visual impairments in pwMS, which are reflected in worse (higher) EDSS.^58,59^ Our study shows, for the first time, greater energetic demand to transition between the motor and visual states compared to HC is associated with disability in MS.

## Limitations

One of the limitations of the study was in the quality of the MRI collected, as the fMRI acquisition time was relatively short and had a longer TR (6 min, TR = 2.3 s) and the dMRI acquisition had only a single b-value of 800 and only 55 directions. A longer fMRI acquisition may result in finding other, less prevalent, dynamic states, while a better dMRI acquisition with greater number of directions may help quantify complex white matter architecture more accurately. Another limitation is the relatively low number of HCs compared to pwMS, which could result in an imbalance of the dynamic states. Lastly, another limitation of the study is the cross-sectional nature of the data; future studies will investigate how brain dynamics change over time and relate to physical and cognitive decline.

## Conclusion

This is the first study using NCT to investigate how dynamic brain states and their associated transition energies change in MS and, furthermore, to associate brain activity entropy with lesion load and NCT-derived brain state transition energies. As hypothesized, we found that global brain activity entropy decreased with increasing lesion load and increasing transition energies. Greater overall transition energies were associated with pwMS who had evidence of disability compared to both HC and pwMS without disability, while decreased overall transition energies were associated with pwMS without disability compared to the other two groups. Transition energies in ventral attention and somatomotor networks largely drove the group classifications. This study, through NCT, sheds light on a possible mechanism of how MS-related damage to the brain’s structural backbone can impact brain activity dynamics, entropy and energetics.

## Acknowledgements

C.T. helped with post-processing, performed the statistical analysis, and wrote the manuscript.

S.C. replicated the analysis using a different brain atlas.

K.J. collected the data, performed pre- and post-processing of MRI data, and reviewed the article.

S.G. collected the data, helped interpret results, and reviewed the article.

A.K. designed and supervised the study, collected the data, and edited the article.

## Ethics statement

All studies were approved by an ethical standards committee on human experimentation and written informed consent was obtained from all patients.

## Abbreviations

ADHD: attention-deficit/ hyperactivity disorder
AMI: adjusted mutual information
AUC: area under ROC curve
BOLD: blood-oxygen level dependent
CER: cerebellum
CIS: clinically isolated syndrome
CNS: central nervous system
DAN: dorsal attention network
DMN: default-mode network
EDSS: expanded disability status scale
fMRI: functional MRI
FP: fronto-parietal
GM: gray matter
HC: healthy controls
LST: lesion segmentation tool
MS: multiple sclerosis
NCT: network control theory
PPMS: primary progressive
MS pwMS: people with multiple sclerosis
SOM: somatomotor
SMOTE: synthetic majority over-sampling technique
SPMS: secondary progressive
SUB: subcortex
VAN: ventral attention
VIS: visual
WM: white matter

