## Supplementary Info for "Larger lesion volume in people with multiple sclerosis is associated with increased transition energies between brain states and decreased entropy of brain activity"

### Lesion volume is related to transition energy and entropy in multiple sclerosis

**Running title:** Network control theory in MS

**Corresponding Author:** Amy Kuceyeski, Ph.D.
**Mailing Address:** 526 Campus Road, Biotechnology building, Room 101D, Ithaca NY 14850

### 1 Assessing the stability of clustering

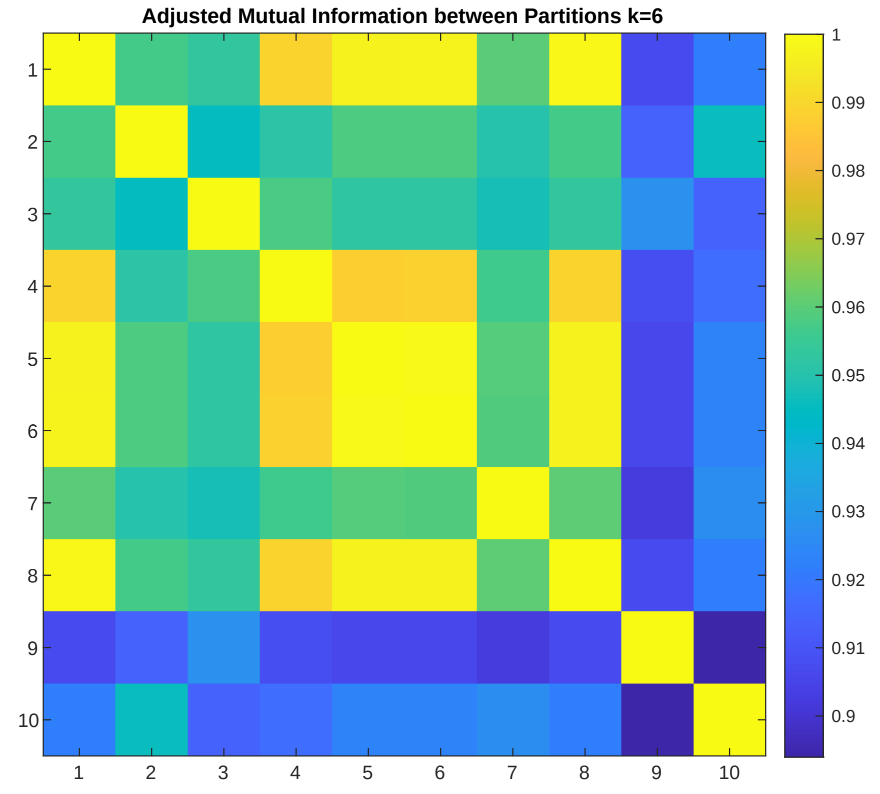


**Figure 1:** The adjusted mutual information shared between 10 independently generated partitions of our data at k = 6. Values range between 0 and 1, where 1 indicates identical partitions.

### 2 The optimal number of clusters

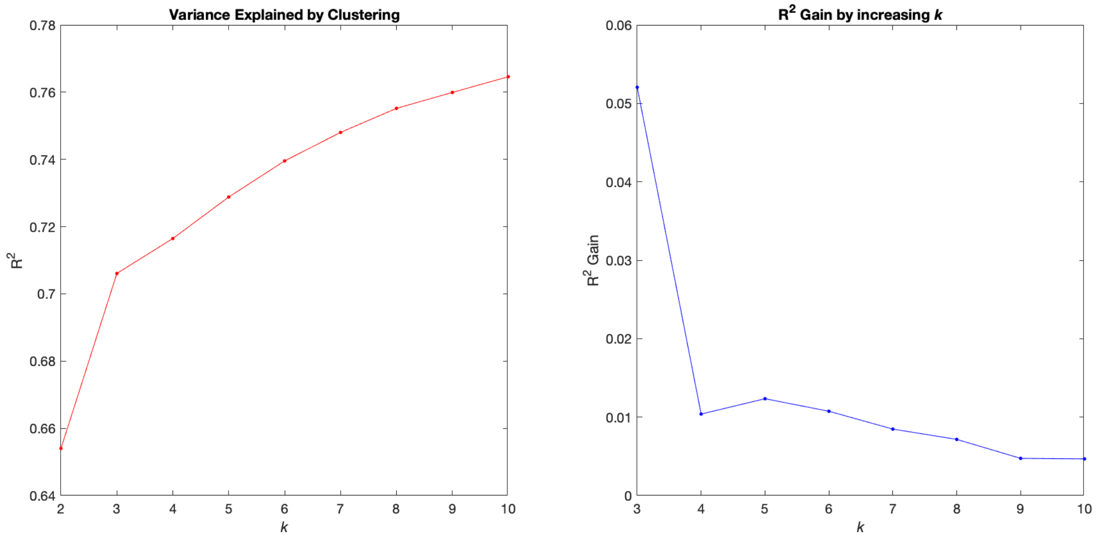


**Figure 2: Choosing the best number of clusters**. Explained variance and gain in explained variance by increasing the number of clusters (k) from 2 to 10. The best number of clusters is k=6 as increasing k beyond k = 6 results in less than a 1% increase of variance explained.

### 3 Comparison of transition probability

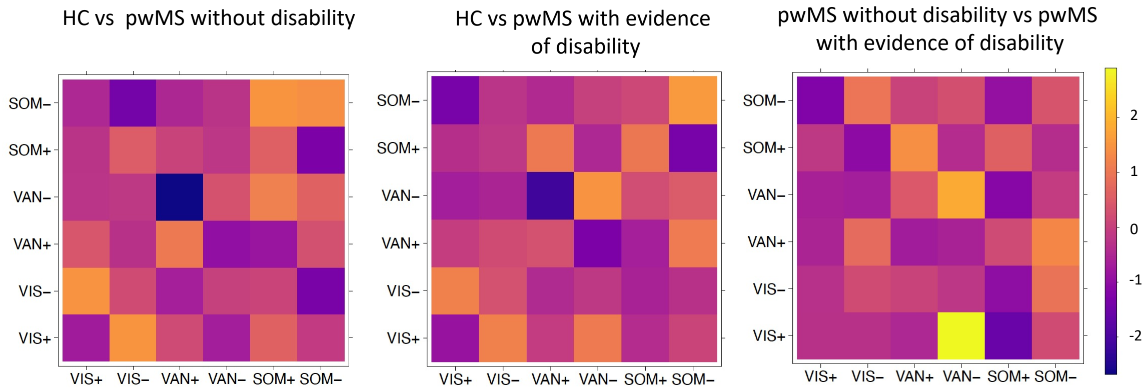


**Figure 3: Comparison of brain state transition probabilities between HC and pwMS disability subgroups for each pairwise transitions.** Group differences are visualized via t-statistic, where the differences were computed as HC - pwMS with no disability, HC - pwMS with evidence of disability, and pwMS with no disability - with evidence of disability. (VIS= Visual, FP= Frontoparietal, SOM= Somatomotor, and VAN= Ventral attention network)

### 4 The metrics of the brain state dynamics

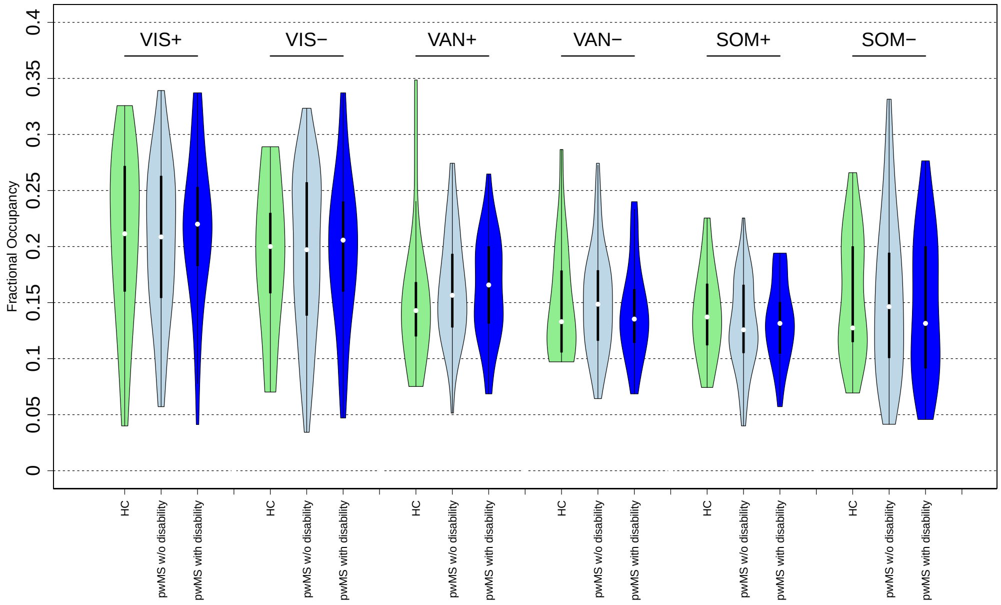


**Figure 4:** Fractional occupancy was presented for each dynamic states obtained for each group (HC, pwMS who had no disability and those who had evidence of disability).

### 5 Optimizing the parameter ”T”

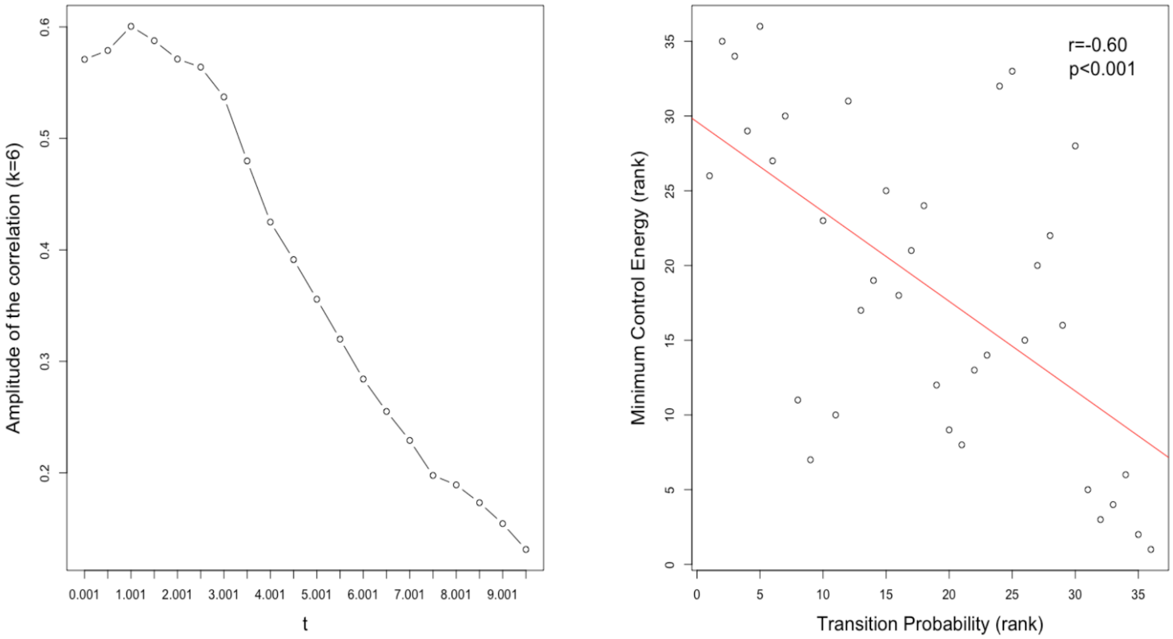


**Figure 5:** Correlation between transition energy and transition probability when varying the parameter T. The highest correlation was obtained when T=1. The figure on the right panel shows the correlation between the rank of transition energy and transition probability using the optimal T=1.

### 6 Replication of the results using cc400

#### 6.1 Finding the best number of clusters

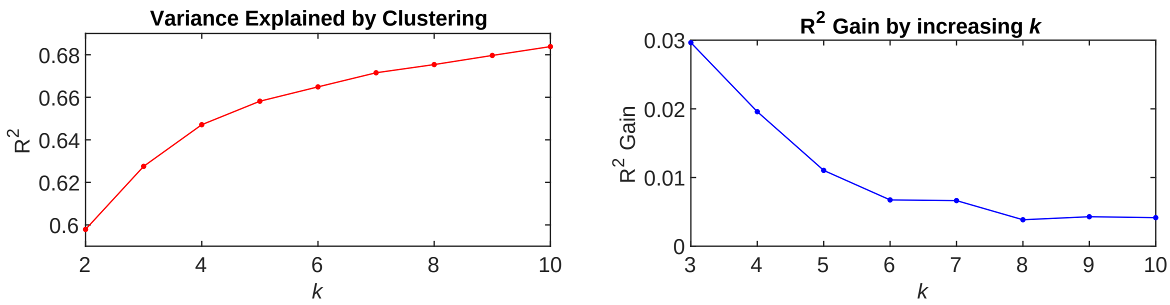


**Figure 6:** Choosing the best number of clusters. Explained variance and gain in explained variance by increasing the number of clusters (k) from 2 to 10. The best number of clusters is k=5 as increasing k beyond k = 5 results in less than a 1% increase of variance explained.

#### 6.2 Centroids with cc400

Figure 7 shows the different centroids and radial plots for each brain state leading to show/label each state and transition.

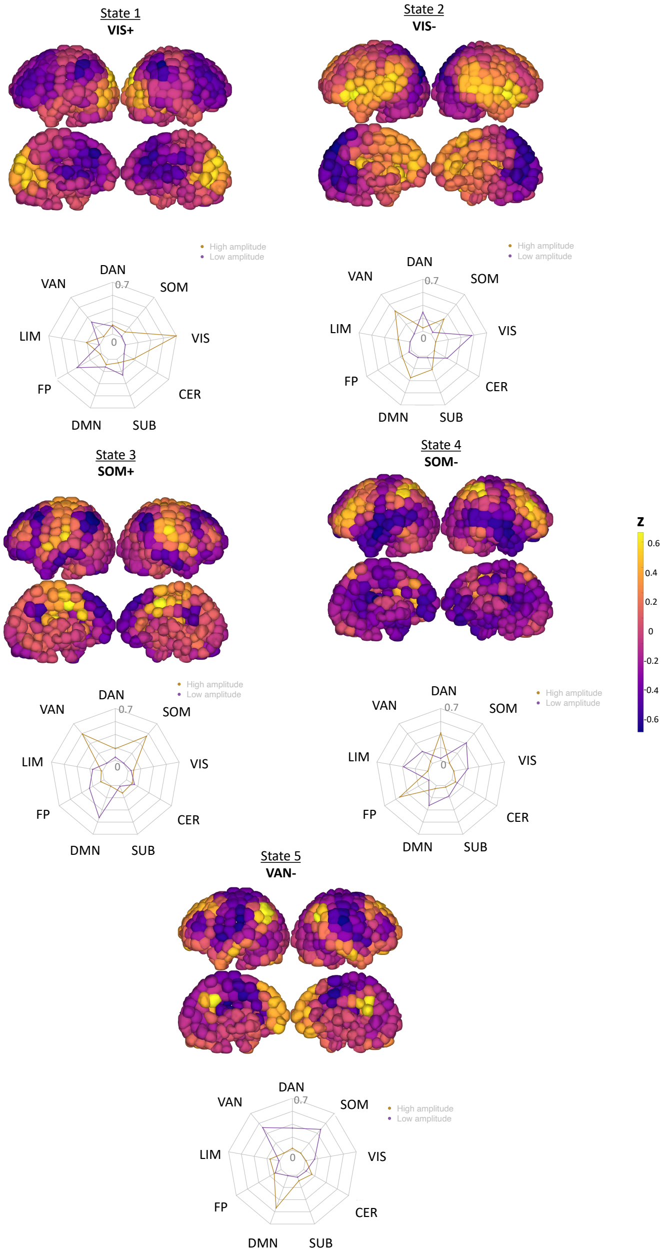


**Figure 7:** The centroid and radial plots for each transition brain state based on the best number of clusters being 5.

#### 6.3 Temporal dynamic metrics with cc400

Figure 8 shows the fractional occupancy obtained using cc400 atlas. These metrics were given for each state in HC, pwMS who had no disability, and those with evidence of disability separately.

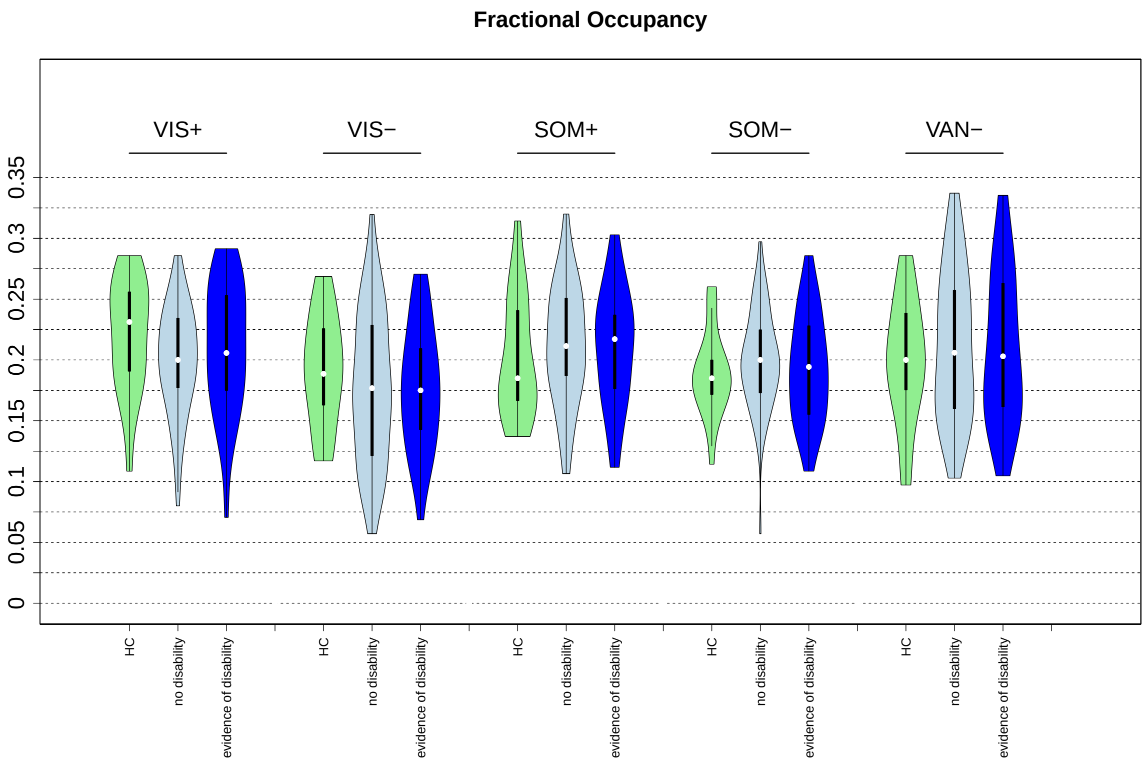


**Figure 8:** The fractional occupancy metrics obtained with cc400 atlas and k=5.

#### 6.4 Comparison of TE obtained with cc400

Figure 9 shows the haufe transformed logistic regression model coefficients for classifying HC vs pwMS w/o disability, HC vs pwMS w/ disability, and pwMS w/o vs pwMS w/ disability using the cc400 atlas.

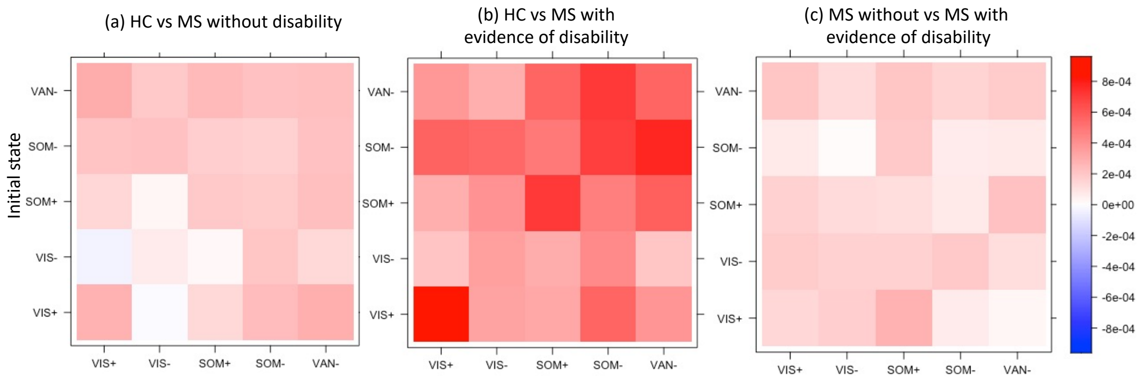


**Figure 9:** Coefficients for the logistic regression with ridge regularization models classifying between HC vs pwMS without disability, HC vs pwMS with disability, and pwMS without disability vs pwMS with disability. (VIS= Visual, VAN= Ventral attention network, and SOM= Somatomotor) *: p-value<0.05 after multiple comparison correction.

#### 6.5 Correlation of entropy with energy

Figure 10 shows the correlation between the average regional entropy vs average transition energy, entropy vs lesion load as well as energy vs lesion load, where energy and entropy were computed using the cc400 atlas. Similar to the results obtained with fs86 atlas, there was a significant negative correlation between entropy and energy (r=-0.23, p-value=0.01).

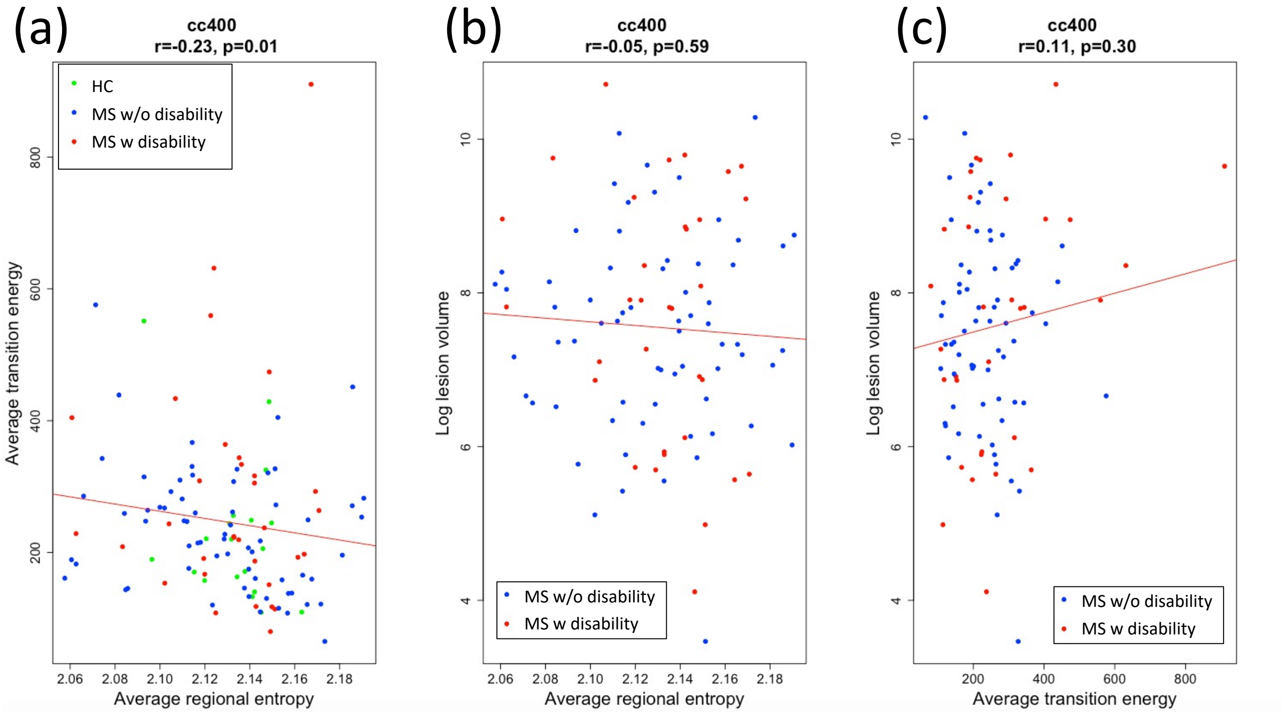


**Figure 10:** The amplitude of the correlation between the (a) energy vs entropy, (b) lesion load vs entropy as well as between (c) lesion load and energy

#### 6.6 Finding best t with cc400

Figure 11 shows the amplitude of the correlation between the rank of the transition probability and rank of the transition energy for different t values between 0.001 and 10. The best value of t was 0.5 with an amplitude of correlation > 0.9.

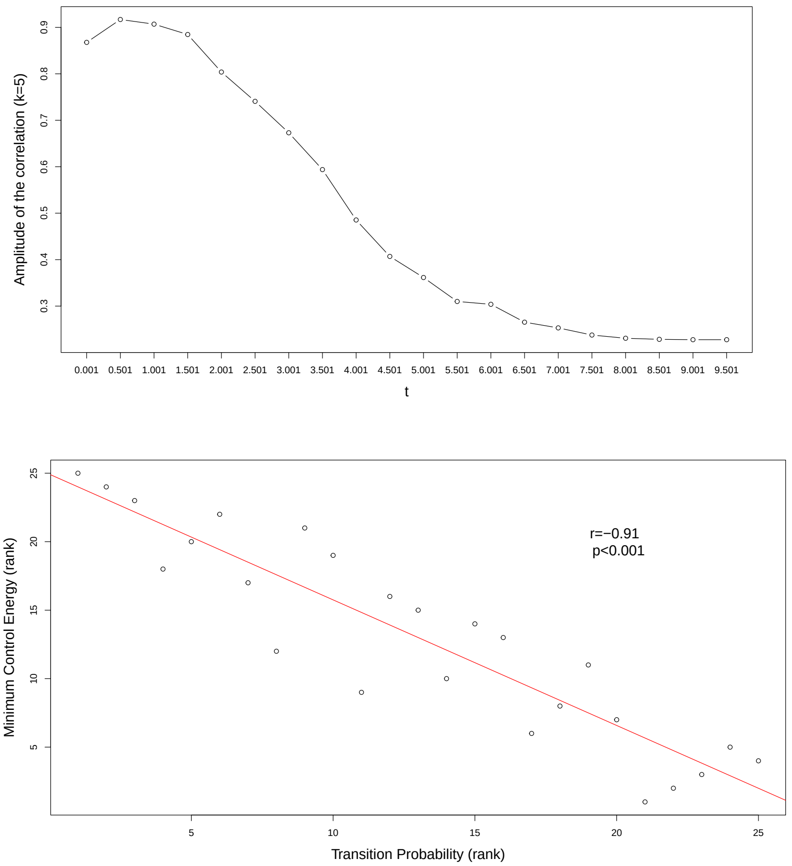


**Figure 11:** The amplitude of the correlation between the rank of the transition probability and rank of the transition energy for varying "t" (top row) and for the best "t" (bottom row)
